# Spontaneous restoration of functional β-cell mass in obese SM/J mice

**DOI:** 10.1101/2020.05.19.104588

**Authors:** Mario A Miranda, Caryn Carson, Celine L St Pierre, Juan F Macias-Velasco, Jing W Hughes, Marcus Kunzmann, Heather Schmidt, Jessica P Wayhart, Heather A Lawson

**Affiliations:** Department of Genetics, Washington University School of Medicine, 660 South Euclid Ave, Saint Louis, MO, USA; Department of Medicine, Washington University School of Medicine, 660 South Euclid Ave, Saint Louis, MO, USA

**Author notes:** Corresponding author 660 South Euclid Ave, Campus Box 8232, Saint Louis, MO, 63110.

**Keywords:** hyperglycemia, insulin, β-cell mass, diabetes, obesity, mouse model

## Abstract

Maintenance of functional β-cell mass is critical to preventing diabetes, but the physiological mechanisms that cause β-cell populations to thrive or fail in the context of obesity are unknown. High fat-fed SM/J mice spontaneously transition from hyperglycemic-obese to normoglycemic-obese with age, providing a unique opportunity to study β-cell adaptation. Here, we characterize insulin homeostasis, islet morphology, and β-cell function during SM/J’s diabetic remission. As they resolve hyperglycemia, obese SM/J mice dramatically increase circulating and pancreatic insulin levels while improving insulin sensitivity. Immunostaining of pancreatic sections reveals that obese SM/J mice selectively increase β-cell mass but not α-cell mass. Obese SM/J mice do not show elevated β-cell mitotic index, but rather elevated α-cell mitotic index. Functional assessment of isolated islets reveals that obese SM/J mice increase glucose stimulated insulin secretion, decrease basal insulin secretion, and increase islet insulin content. These results establish that β-cell mass expansion and improved β-cell function underlie the resolution of hyperglycemia, indicating that obese SM/J mice are a valuable tool for exploring how functional β-cell mass can be recovered in the context of obesity.

## Introduction

Obesity and diabetes are a deadly combination, compounding risk for cardiovascular disease, cancer, and stroke(30, 65, 95). Obesity raises the risk of developing type 2 diabetes 27-76 fold, while approximately 60% of individuals with diabetes are obese(1, 12, 15, 19). Chronic obesity exerts glycemic stress on pancreatic β-cells, causing dysregulation and dysfunction, ultimately resulting in hyperglycemia(49, 67, 77, 86). Despite the stress obesity places on β-cells, 10-30% of obese individuals maintain glycemic control and are at low risk for developing diabetes(61). These low-risk obese individuals have elevated β-cell mass and improved insulin secretion compared to BMI-matched diabetic-obese individuals(2, 9, 75, 90). Understanding the differences in β-cell physiology between these populations may reveal therapeutic strategies for maintaining and improving glycemic control in obese individuals.

Recent work suggests β-cells do not respond uniformly to glycemic stress, rather they experience variable fates including dedifferentiation, replication, and apoptosis(10, 18, 34). Understanding how these changes mediate diabetic risk is complicated by β-cells heterogeneity. β-cell populations include subtypes that specialize in basal insulin secretion, β-cell replication, coordinating “hub” cells, and β-cells derived from transdifferentiated α-cells, each of which differ in glycemic stress response(31, 42, 81, 88). Thus, determining what differentiates nondiabetic-obese and diabetic-obese populations requires connecting β-cell subtypes to their fate in prolonged glycemic stress. Like in humans, diabetic risk in obese mice depends on genetic background(44, 48, 80). Variation in β-cell heterogeneity likely underlies variability in islet stress response, and thus needs to be accounted for when comparing nondiabetic-obese and diabetic-obese populations. Loss of function mutations in leptin (*ob/ob*) and leptin receptor (*db/db*) provide insight into β-cell physiology in nondiabetic-obese and diabetic-obese states within individual mouse strains(8, 40, 46, 53), however leptin and its receptor play a critical role in β-cell function independent of obesity, limiting interpretations of these studies(22). No current mouse model is well-suited to examine physiological differences in β-cell health between nondiabetic-obese and diabetic-obese states.

The SM/J inbred mouse strain has traditionally been used to study interactions between diet and metabolism, and more recently has uncovered genetic architecture underlying diet-induced obesity and glucose homeostasis(17, 49–52, 63). After 20 weeks on a high fat diet, SM/J mice display characteristics of diabetic-obese mice, including elevated adiposity, hyperglycemia, and glucose intolerance(27). We have previously shown that by 30 weeks of age, high fat-fed SM/J mice enter diabetic remission, characterized by normalized fasting blood glucose, and greatly improved glucose tolerance and insulin sensitivity(11). Importantly, these changes occur in the context of sustained obesity. Given the central role of β-cell health in susceptibility to diabetic-obesity, we hypothesize that obese SM/J mice undergo restoration of functional β-cell mass during the resolution of hyperglycemia. This study focuses on how insulin homeostasis, β-cell morphology, and β-cell function change during this remarkable transition and establishes SM/J mice as a useful model for teasing apart diabetic-obese and nondiabetic-obese states.

## Methods

### Animal husbandry and tissue collection

SM/J mice were obtained from The Jackson Laboratory (Bar Harbor, ME). Experimental animals were generated at the Washington University School of Medicine and all experiments were approved by the Institutional Animal Care and Use Committee in accordance with the National Institutes of Health guidelines for the care and use of laboratory animals. Mice were weaned onto a high fat diet (42% kcal from fat; Envigo Teklad TD88137) or an isocaloric low fat diet (15% kcal from fat; Research Diets D12284), as previously described(11). At 20 or 30 weeks of age, mice were fasted for 4 hours, and blood glucose was measured via glucometer (GLUCOCARD). Mice were then injected with an overdose of sodium pentobarbital, followed by a toe pinch to ensure unconsciousness. Blood was collected via cardiac puncture and pancreas was detached from the spleen and duodenum.

### Serum and pancreatic insulin measurements

Blood obtained via cardiac puncture was spun at 6000 rpm at 4°C for 20 minutes to separate plasma, which was collected and stored at −80 °C. Whole pancreas was homogenized in acid ethanol and incubated at 4°C for 48 hours, shaking. Homogenate was centrifuged at 2500 rpm for 30 min at 4°C. Supernatant was collected and stored at −20°C. Protein content was measured using Pierce BCA Protein Assay kit (Thermo Scientific) according to manufacturer’s instructions and read at 562 nm on the Synergy H1 Microplate Reader (Biotek). Insulin ELISA (ALPCO 80-INSMR-CH01) was used to measure plasma and pancreatic insulin levels following manufacturer’s instructions.

### Insulin Tolerance Test

At 19 or 29 weeks of age, mice were fasted for 4 hours prior to procedure. Insulin (humulin) was prepared by mixing 10 ul insulin with 10 ml sterile saline. Mice were injected with 3.75 ul insulin mixture/g bodyweight. Blood glucose levels were assessed from a tail nick at times = 0, 15, 30, 60, and 120 minutes via glucometer (GLUCOCARD).

### Islet Histology and Analyses

At the time of tissue collection, whole pancreas was placed in 3 mL of neutral buffered formalin. These samples were incubated at 4°C while gently shaking for 24 hours. Immediately afterwards, samples were placed into plastic cages and acclimated to 50% EtOH for 1 hour. Samples were then processed into paraffin blocks using a Leica tissue processor with the following protocol: 70% EtOH for 1 hour x 2, 85% EtOH for 1 hour, 95% EtOH for 1 hour x 2, 100% EtOH for 1 hour x 2, Xylenes for 1 hour x 2, paraffin wax. Pancreas blocks were sectioned into 4 μm thick sections. Four samples per individual were randomly selected, at least 100 μm apart.

Slides were incubated at 60°C for 1 hour, then placed in xylenes to remove remaining paraffin wax. Slides were then rehydrated using successive decreasing EtOH concentrations (xylenes x 2, 50% EtOH in xylenes, 100% EtOH x 2, 95% EtOH, 70% EtOH, 50% EtOH, H2O). Slides were incubated in sodium citrate (pH 6) at 85°C for 30 minutes, then submerged in running water for 5 minutes. Slides were washed with 0.025% Triton X-100 in TBS and blocked in 10% normal donkey serum for 1 hour (Abcam ab7475), followed by incubation with primary antibody overnight at 4°C. [Primary antibodies: rat anti-insulin (1:100, R&D MAB1417), mouse anti-glucagon (1:100, Abcam ab10988), and rabbit anti-phospho-histone H3 (1:100, Sigma SAB4504429)]. After an additional wash, secondary antibody was applied for 1 hour at room temperature. [Secondary antibodies: donkey anti-rabbit 488 (1:1000, Abcam ab150061), donkey anti-mouse 647 (1:1000, Abcam ab150107), and donkey anti-rat 555 (1:1000, Abcam ab 150154)]. Fluoroshield Mounting Medium with DAPI (Abcam) was applied to seal the coverslip and slides were stored at 4°C. Imaging was performed using the Zeiss AxioScan. Z1 at 20X magnification and 94.79% laser intensity.

Background was subtracted from DAPI, insulin, glucagon, and phospho-histone H3 images using ImageJ. DAPI channel was used to identify total nuclei in CellProfiler. Insulin and glucagon channels were combined and overlaid on the DAPI image to identify islet nuclei. Insulin (INS^+^) staining overlaid with DAPI identified β-cell cells, glucagon (GCG^+^) staining overlaid with DAPI identified α-cells. Phosphohistone H3 (PHH3^+^) staining identified mitotic nuclei. Total nuclei, islet cells, β-cells, α-cells, and mitotic nuclei were summed across 4 slides for each individual. Islet, β-cell, and α-cell mass is reported as fraction of total nuclei. Mitotic islet index is reported as proportion of β-cells and α-cells positive for phosphohistone H3. Islets with diameter < 50 μm were discarded.

### Islet isolation

Pancreas was removed and placed in 8mL HBSS buffer on ice. Pancreas was then thoroughly minced. Collagenase P (Roche) was added to a final concentration of 0.75 mg/ml. Mixture was then shaken in a 37°C water bath for 10-14 minutes. Mixture was spun at 1500 rpm for 2 minutes. The pellet was washed twice with HBSS. The pellet was re-suspended in HBSS and transferred a petri dish. Hand-selected islets were placed in sterile-filtered RPMI with L-glutamine (Gibco) containing 11mM glucose, supplemented with 5% pen/strep and 10% Fetal Bovine Serum (Gibco). Islets were rested overnight in a cell culture incubator set to 37°C with 5% CO_2_.

### Glucose Stimulated Insulin Secretion and Islet Insulin Content

Islets of roughly equal size were equilibrated in KRBH buffer containing 2.8 mM glucose for 30 minutes at 37°C. 5 islets were hand selected and placed in 150 μl KRBH containing either 2.8 or 11 mM glucose. Tubes were placed in a 37°C water bath for 45 min. Islets were then spun at 2000 x g, hand-picked with a pipette, and transferred from the secretion tube and placed in the content tube with acid ethanol. The content and secretion tubes were stored at −20°C overnight. Each condition was performed in duplicate for each individual. Mouse insulin ELISA (ALPCO 80-INSMU-E01) was performed according to manufacturer’s instructions, with the secretion tubes diluted 1:5, and content tubes diluted 1:100. Normalized insulin secretion was calculated by dividing the secreted value by the content value. Glucose stimulated insulin secretion was calculated by dividing the normalized insulin secretion at 11mM glucose by the normalized insulin secretion at 2.8 mM glucose. Each sample was measured in duplicate. Total islet protein within each content tube was measured using Pierce BCA Protein Assay kit (Thermo Scientific) according to manufacturer’s instructions and read at 562 nm on the Synergy H1 Microplate Reader (Biotek). Islet insulin content was calculated by dividing the insulin level in the content tubes by the total protein value.

### Statistical analyses

Phenotypes were assessed for normality by a Shapiro-Wilk test, and outliers removed. A student’s t-test was used to assess significance between two cohorts, while a one-way ANOVA with Tukey’s Post Hoc test was used to assess significance among multiple cohorts. Pearson’s correlation was used to determine strength of correlation among variables. P-values < 0.05 were considered significant.

## Results

### Obese SM/J mice increase insulin levels and improve insulin sensitivity

The resiliency of β-cells distinguishes nondiabetic-obese and diabetic-obese individuals(8, 46, 47, 63, 66, 72, 74, 81). While both groups develop hyperinsulinemia, diabetic-obese individuals become insulin resistant, leading to β-cell dysfunction, hypoinsulinemia, and hyperglycemia. Our previous work shows that obese SM/J mice spontaneously transition from hyperglycemic to normoglycemic with age(11). Principle to this is a 40 mg/dl decrease in fasting glucose levels in high fat-fed SM/J mice between 20 and 30-weeks (Fig. 1A). We first sought to characterize how insulin homeostasis changes during this transition. Interestingly, 20-week high fat-fed SM/J mice have comparable levels of plasma and pancreatic insulin levels compared to age-matched low fat-fed mice (Fig. 1B-C). By 30 weeks, high fat-fed SM/J mice increase circulating insulin levels 5.3-fold and pancreatic insulin levels 1.9-fold, in line with other models of hyperinsulinimic nondiabetic-obesity(33, 36, 55). We sought to test for peripheral insulin resistance via an insulin tolerance test (ITT), as insulin resistance is a known mechanism for increasing circulating and pancreatic insulin levels. Surprisingly, 20-week high fat-fed SM/J mice display insulin resistance compared to low fat-fed mice, however, insulin sensitivity is restored by 30 weeks (Fig. 1D-E). The simultaneous increase in insulin production and improved insulin sensitivity is unprecedented and suggests a novel mechanism beyond insulin resistance for enhancing β-cell insulin secretion.

**Figure 1.**
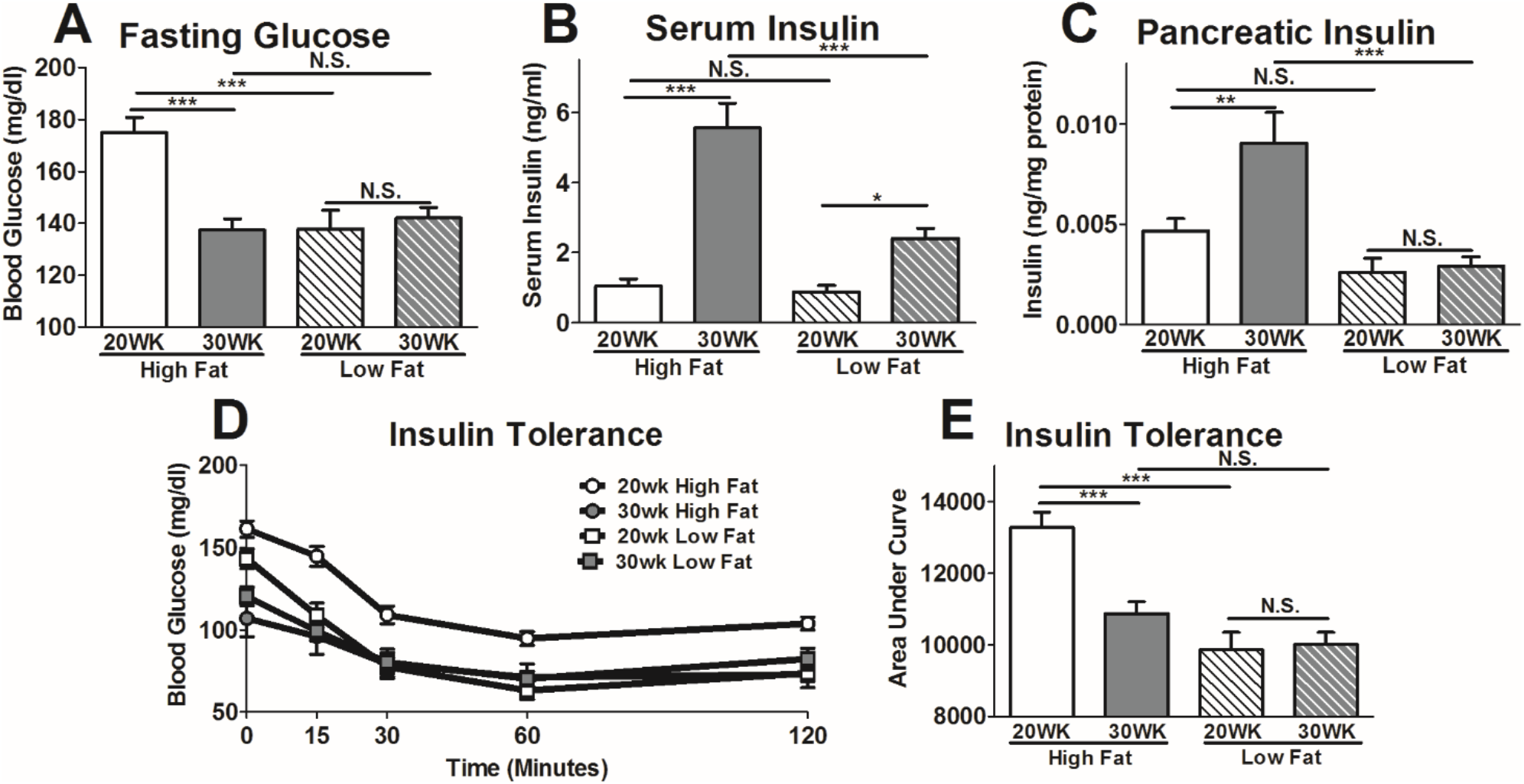
Insulin homeostasis during the resolution of hyperglycemia in obese SM/J mice. Blood glucose levels measured after 4-hour fast from high and low fat-fed, 20- and 30-week SM/J mice (**A)**. Plasma insulin (**B)**and pancreatic insulin levels (**C)**assessed via insulin ELISA, collected after 4-hour fast. Insulin tolerance test performed via intraperitoneal insulin injection following 4-hour fast (**D)**, summarized in the area under the curve (**E)**. N = 38-50 for panel A, C, D. N = 10-24 for panel B-C, equal numbers of males and females. Bar represents group means, error bars represent SEM. *p<0.05, **p<0.01, ***p<0.001, N.S. Not Significant.

### Obese SM/J mice increase islet mass during resolution of hyperglycemia

In humans and mice, obesity initially increases islet mass, and maintenance of that mass in part differentiates nondiabetic-obese individuals from diabetic-obese individuals(2, 9, 25, 59, 76, 85). To understand the source of increased insulin production in obese SM/J mice, we examined islet morphology during the resolution of hyperglycemia. To quantify islet mass and number, β-cell mass, α-cell mass, and mitotic index, we randomly selected 4 sections per fixed pancreas and stained with antibodies against insulin, glucagon, and phospho-histone H3. Representative images of immuno-stained pancreatic sections for 30-week high fat-fed mice and 30-week low fat-fed mice are shown in Figure 2A-B. Consistent with other mouse models of obesity, 20-week high fat-fed SM/J mice have a 2.75-fold increase in total islet mass compared to low fat-fed mice (Fig. 2C). This increased mass is driven by an increase in both median islet area and number of islets (Fig. 2D-E). Islet mass is further elevated 2-fold between 20- and 30-weeks in high fat-fed mice, while the islet population remains unchanged in low fat-fed mice. A full summary of the islet quantification is presented in Supplemental Table 1. Distribution of islet size is shown in Supplemental Figure 1, along with corresponding density plot for each cohort. Islet mass correlates with BMI in obese humans(26), a similar correlation is seen between islet mass and body weight in high fat-fed SM/J mice (Fig. 2F).

**Figure 2.**
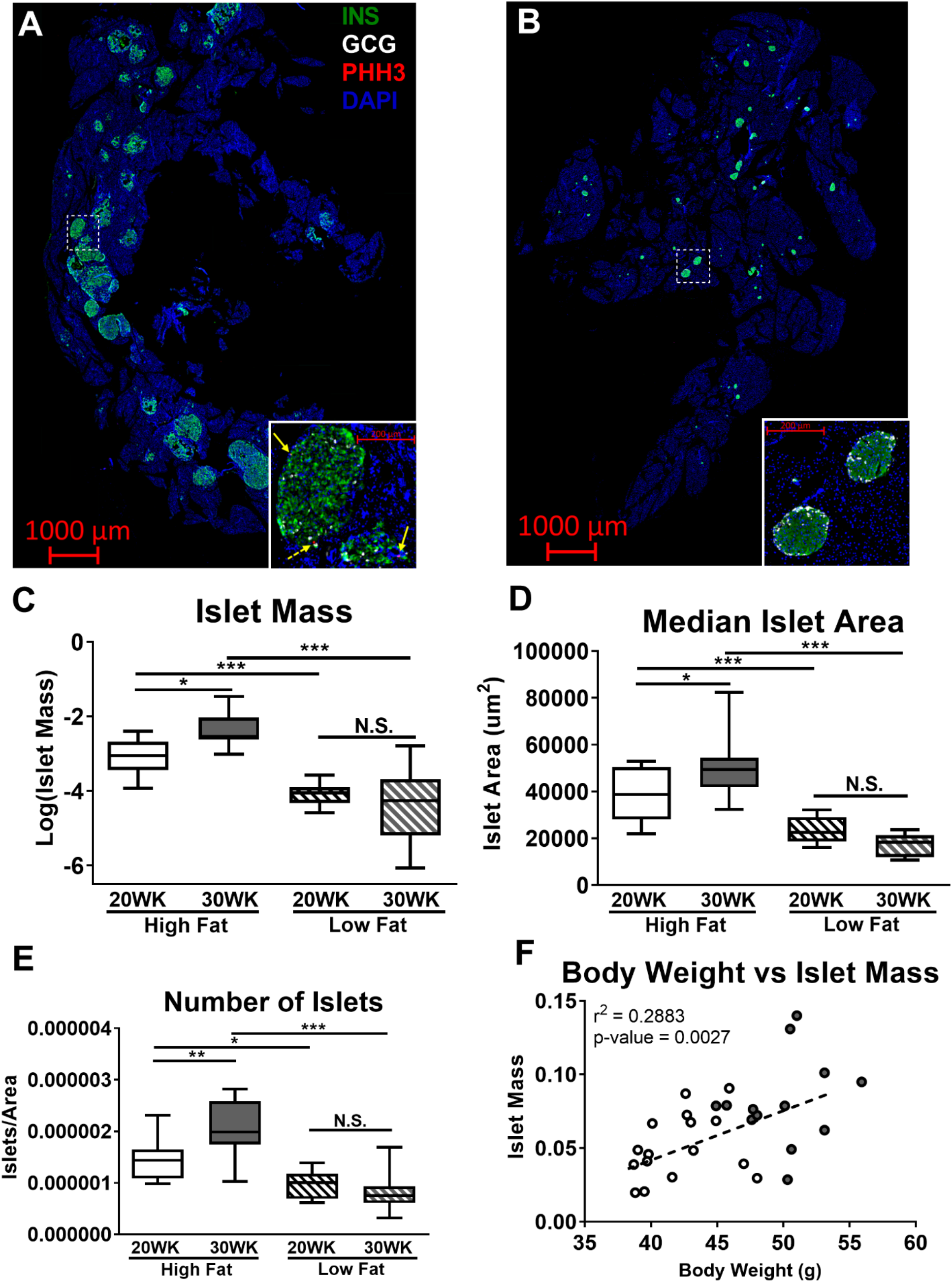
Changes in islet mass during the resolution of hyperglycemia. Representative pancreatic cross sections from 30-week high fat-fed mice **(A)**and 30-week low fat-fed mice **(B)**stained for insulin (green), glucagon (white), and phosphohistone H3 (red). Dashed white box identifies location of image in inset. Solid yellow arrows within inset identify INS^+^:PHH3^+^ cells, dashed yellow arrow identifies GCG^+^:PHH3^+^ cell. Islet mass reported as ratio of islet cells over total cells, summed across 4 pancreatic sections **(C)**. Median islet area calculated for each individual across 4 sections **(D)**. Total number of islets quantified per individual, normalized by total DAPI area **(E)**. Correlation between body weight and β-cell mass in high fat-fed mice **(F)**, open circles – 20-week high fat-fed, filled circles – 30-week high fat-fed. N = 12-16 per cohort, equal number of males and females. *p<0.05, **p<0.01, ***p<0.001, N.S. Not Significant.

### Obese SM/J mice increase β-cell mass and α-cell replication

To identify the source of the increased islet mass in high fat-fed SM/J mice, we quantified β-cell and α-cell mass within each cohort. Increased islet mass in 20-week high fat-fed mice is driven by a 3.3-fold increase in β-cell mass and a 2.5-fold increase in α-cell mass compared to low fat mice, while growth between 20- and 30-week high fat-fed mice is driven by a further 2.2-fold increase in β-cell mass (Fig. 3A-B). In obesity, islet mass expands primarily through β-cell replication (34, 68, 85, 92). We quantified mitotic index of β- and α-cells in our model using phosphohistone H3 and assessed how mitotic index relates to β-cell mass during the resolution of hyperglycemia in obese SM/J mice. Surprisingly, calculation of β-cell mitotic index reveals similar rates of β-cell replication across cohorts (Fig. 3C), while α-cell mitotic index is elevated 6-fold in high fat-fed mice compared to low fat-fed controls (Fig. 3D). Examining the relationship between β-cell mitotic index and β-cell mass in high fat-fed mice reveals β-cell replication correlates with β-cell mass in 20-week mice, but not 30-week mice (Fig. 3E-F).

**Figure 3.**
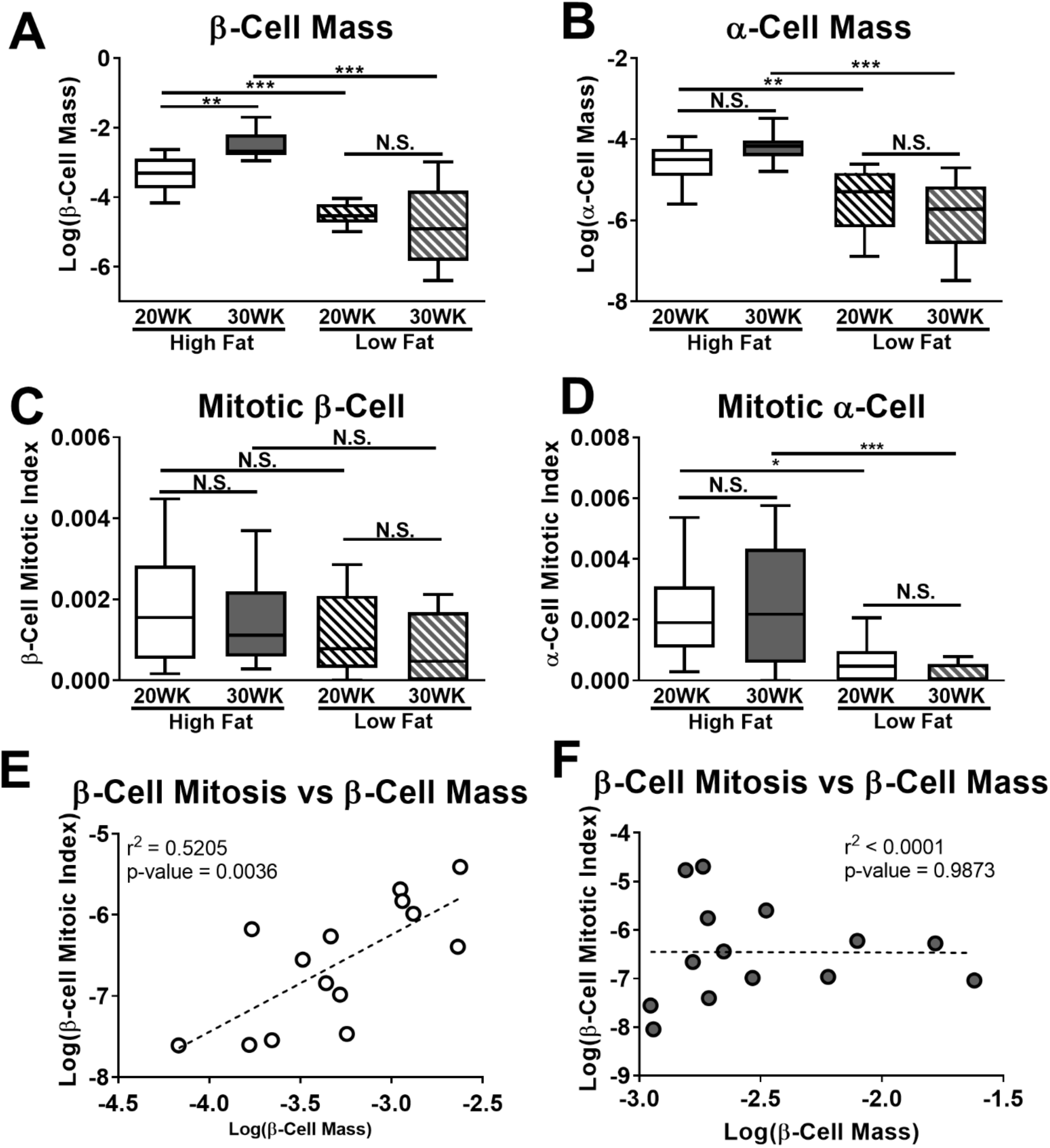
Islet cell mass and mitotic index in obese SM/J mice. β-cell mass reported as ratio of INS^+^ cells divided by total cells summed across 4 slides per individual (**A)**. α-cell mass reported as GCG^+^ cells divided by total cells summed across 4 slides per individual (**B)**. β-cell mitotic index calculated by dividing INS^+^:PHH3^+^ cells divided by total INS^+^ cells summed across 4 slides per individual (**C)**. α-cell mitotic index calculated by dividing GCG^+^:PHH3^+^ cells by total GCG^+^ cells summed across 4 slides (**D).**Correlation between β-cell mitotic index and β-cell mass in 20 week high fat-fed mice (**E)**and 30-week high fat-fed mice (**F)**. Open circles – 20-week high fat-fed, filled circles – 30-week high fat-fed. N = 12-16 per cohort, equal males and females. *p<0.05, **p<0.01, ***p<0.001, N.S. Not Significant.

### Obese SM/J mice increase islet insulin secretion and insulin content

In conjunction with changing β-cell morphology, diabetic-obesity is associated with altered β-cell function, including diminished first phase insulin secretion, increased basal insulin secretion, and decreased β-cell insulin production (16, 23, 57, 66). We sought to examine if changes in β-cell insulin secretion and content corresponded with the resolution of hyperglycemia and expanded β-cell mass we observe. To test this, we isolated islets from high and low fat-fed 20- and 30-week SM/J mice. After allowing islets to rest overnight, we performed a glucose-stimulated insulin secretion assay by subjecting islets to low (2.8 mM) or high (11 mM) glucose conditions. We find that high fat-fed SM/J mice dramatically improve glucose-stimulated insulin secretion between 20 and 30 weeks of age. This includes transitioning from blunted insulin secretion under high glucose conditions to appropriately elevated secretion (Fig. 4A), and improvement in the ratio of insulin secreted in response to high vs low glucose conditions (Fig. 4B). 20-week high fat-fed mice have elevated insulin secretion in response to low glucose (Fig. 4C), consistent with other studies of islets in type 2 diabetic humans and mice. Correspondingly, 20-week high fat-fed SM/J mice have decreased islet insulin content (Fig. 4D), which increases 3-fold by 30 weeks. Consistent with current understanding of the β-cell maturation process(76), there is a positive correlation between obese SM/J islet insulin content and glucose-stimulated insulin secretion (Fig. 4E). This suggests that obese SM/J mice experience β-cell maturation between 20 and 30 weeks, characterized by increased insulin content and improved insulin secretion in response to high glucose. This spontaneous improvement in β-cell health and function in the context of obesity has not been reported in other mouse strains, suggesting a genetic basis unique to SM/J.

**Figure 4.**
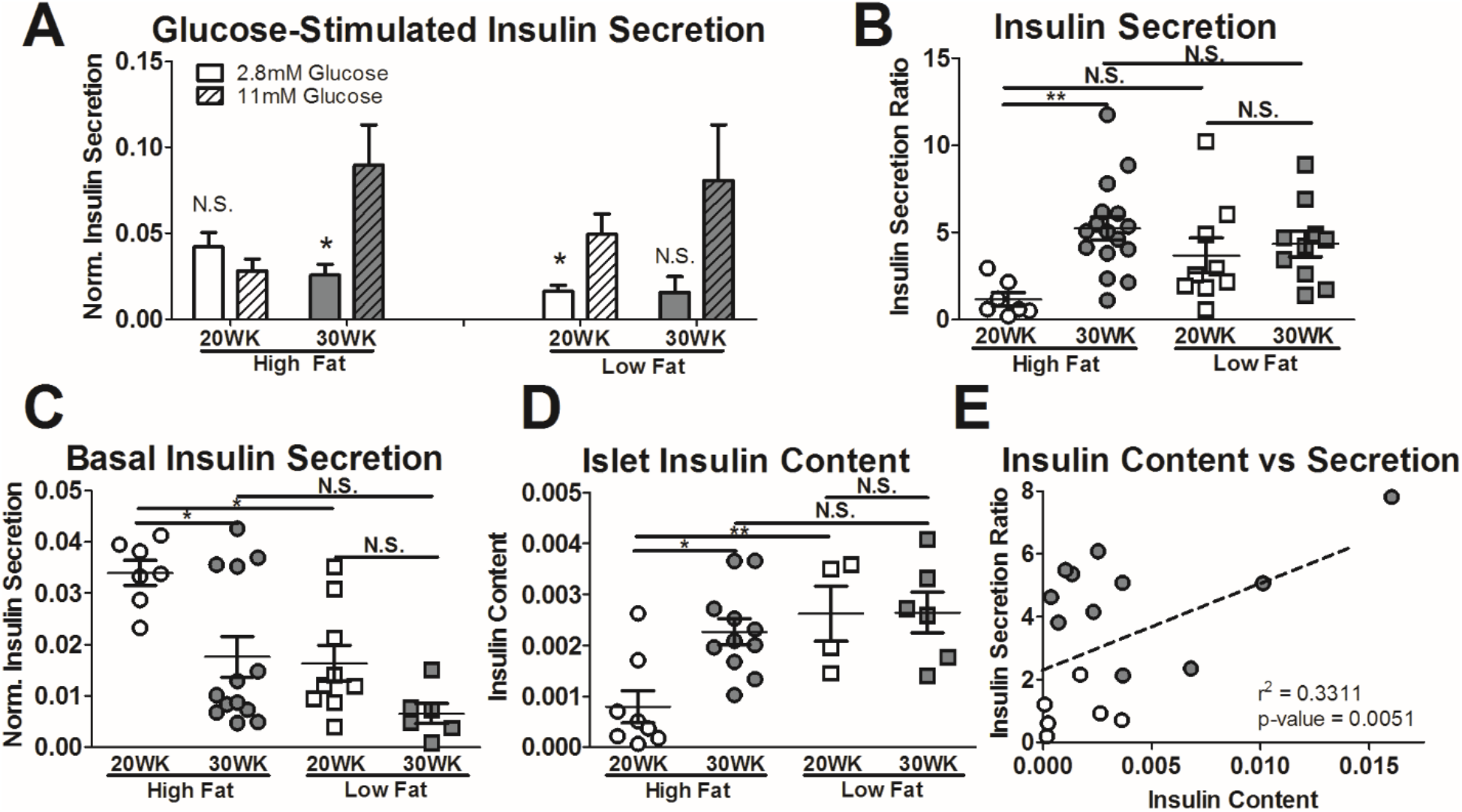
Islet insulin secretion and insulin content. Islet insulin secretion in response to low (2.8 mM) and high (11mM) glucose conditions, normalized by islet insulin content (**A)**, reported as a ratio of high glucose to low glucose insulin secretion (**B)**. Comparison of islet insulin secretion under low glucose conditions in 20- and 30-week, high and low fat-fed mice (**C)**. Islet insulin content normalized by total protein measured via protein BCA (**D)**. Correlation between insulin secretion ratio and islet insulin content (**E)**. Open circles – 20-week high fat-fed, closed circles – 30-week high fat-fed. *p<0.05, **p<0.01, ***p<0.001, N.S. Not Significant.

## Discussion

The ability to maintain appropriate insulin production and secretion, termed functional β-cell mass, is a central determinant of diabetic risk. In this study, we describe insulin homeostasis, islet morphology, and β-cell function in obese SM/J mice as they transition from hyperglycemic to normoglycemic. We determine that increased insulin production and insulin sensitivity accompany improved glycemic control, driven by expanded β-cell mass and improved glucose-stimulated insulin secretion. Our results show obese SM/J mice undergo restoration of functional β-cell mass, providing an opportunity to explore how compensatory insulin production can be achieved in the context of obesity.

Susceptibility to high fat diet-induced diabetes in mice depends on genetic background. Several strains and sub-strains develop diabetic-obesity, including hyperglycemia, glucose intolerance, and insulin resistance, consistent with the diabetic phenotypes observed in obese SM/J mice at 20 weeks (3, 44, 83). Remarkably, by 30 weeks, obese SM/J mice have characteristics of diabetic-resistant obese strains, retaining glycemic control by dramatically increasing insulin production and improving insulin sensitivity (3, 79, 83). To our knowledge, this is the first report of transient hyperglycemia in an inbred strain, although similar phenomena have been reported in mice with the leptin receptor (*db/db*) mutation. C57bl/6J *^(db/db)^* and 129/J *^(db/db)^* mice are obese and initially develop mild hyperglycemia at 8-10 weeks of age, but this resolves by 20-30 weeks, concurrent with increased insulin production and β-cell mass(40, 54). Unfortunately, leptin and its receptor play an important role in β-cell growth and function independent of obesity, which confounds understanding of how genetic background mediates diabetic risk in obesity(22).

High fat diet-induced obesity in mice can result in increased islet mass, no change, or decreased mass (3, 39, 66, 79). Across these studies, inability to expand islet mass is associated with hyperglycemia. In humans, islet mass correlates with BMI in nondiabetic obese-individuals, while diabetic-obese individuals have low islet mass compared to nondiabetic individuals (26, 47, 54). High fat-fed SM/J mice are unique because they have expanded islet mass at 20 weeks, yet normal insulin levels and insulin resistance. By 30-weeks, islet mass continues to expand, driven by increased islet area and increased islet number, corresponding with increased insulin production and improved insulin sensitivity. Islet neogenesis may contribute to the increased islet number, and fission of large islets has been reported during development, suggesting islets have mechanisms to maintain an appropriate size(41, 80).

β-cell expansion is the primary driver of islet expansion in mouse models of obesity(8, 46). Some nondiabetic obese mice experience increased β-cell mass, but do not show evidence for elevated β-cell replication in immunostaining of pancreatic sections(38, 83). This has been attributed to islets in the tail of the pancreas being substantially more proliferative in response to high fat diet than the body and head regions(28), thus technical artifacts in sampling could result in inflated variances which mask biological differences. This is could be the case here, given that high fat-fed SM/J’s β-cell mass is far above low fat-fed controls, that their β-cell mass expands 2-fold during the resolution of hyperglycemia, yet we find no evidence for increased β-cell replication. However, α-cell mass also expands in obesity (29, 37, 61). While α-cell mass is elevated in high fat-fed SM/J mice compared to low fat-fed controls, we find it does not change between 20 and 30 weeks, despite substantial elevation of α-cell mitotic index.

Retention of β-cell function separates diabetic-obesity and nondiabetic obesity (5, 35, 45). 20-week high fat-fed SM/J mice have an insulin secretion profile similar to diabetic-obese mice and humans, including blunted glucose-stimulate insulin release, elevated basal insulin secretion, and low islet insulin content, which resolves by 30 weeks. Underscoring this transition is the positive correlation between glucose-stimulated insulin release and islet insulin content. Care was taken to select normal sized islets across all cohorts for functional assessment (~100μm in diameter) indicating this robust improvement in β-cell functional mass is due to changes to β-cell physiology.

Three current, non-mutually exclusive components of β-cell stress response may shed light on the perplexing improvement in glycemic control seen in SM/J mice: β-cell dedifferentiation, nascent β-cell maturation, and changes in β-cell subtype proportions. While early studies concluded overworked β-cells undergo apoptosis (10, 56, 67, 73), recent studies have suggested β-cells dedifferentiate into a dysfunctional, progenitor-like state, potentially as a defense mechanisms against prolonged glycemic stress (18, 43, 58, 84). These dedifferentiated β-cells have low insulin content and poor glucose-stimulated insulin secretion. Further, the dedifferentiated state is reversible in cultured conditions, revealing potential for therapeutic intervention(24). It is feasible that obese SM/J mice have β-cells in the dedifferentiated state at 20-weeks, which would explain the low insulin content and poor functionality despite the elevated β-cell mass. Improvement in insulin sensitivity could ease glycemic stress, allowing dedifferentiated β-cells to redifferentiate by 30 weeks, reestablishing insulin production and secretion.

Work from several groups suggests β-cells can be divided into two broad categories: functionally immature and functionally mature cells. Immature β-cells have greater proliferative potential and are resistant to stress, at the expense of functional maturity (4, 7, 69, 89). These immature β-cells have low insulin content, high basal insulin secretion, and a lack of glucose stimulated insulin secretion. The large β-cell expansion seen in obese SM/J mice, suggests nascent β-cells must undergo maturation at some point. We have no evidence of enhanced β-cell replication at 20-weeks, but it is possible these β-cell are still functionally immature and reach maturity by 30-weeks. This could explain why islets from these mice lack glucose stimulated insulin release, show elevated basal insulin secretion, and have low insulin content, despite elevated mass.

Recent advances in single cell technology has allowed for identification of β-cell subtypes, based on functional characteristics and gene expression. These include β-cells that specializes in basal insulin secretion, characterized by low mature insulin content, and enriched in *db*/*db* diabetic islets(32). While these cells are not equipped to respond to elevated glucose, they are enriched for maturity markers including *Ucn3* and *Glut2*, distinguishing them from immature β-cells. Pancreatic multipotent progenitors (PMPs) are rare insulin positive cells capable of generating endocrine cells *in vivo* including functionally mature β-cells(73, 85). These cells resemble immature β-cells, with low insulin content and *Glut2* expression, whose proliferation is stimulated by glycemic stress in STZ-treated and NOD mouse models. Lastly, β-cell hub cells coordinate calcium signaling and insulin release of surrounding β-cells(42). These cells have markers for both mature and immature β-cells, including expression of *Gck* and *Pdx1,* but low insulin content, and are especially sensitive to glycemic and inflammatory stress. Ablation of these cells results in loss of coordinated insulin release, suggesting they are necessary for mature islet function. Given the elevated β-cell mass, poor insulin secretion, and low insulin content in 20-week high fat-fed SM/J mice, it is possible islets are enriched for basal insulin secretors and PMP’s, while devoid of hub cells. At 30 weeks, as glycemic stress diminishes, basal insulin secretors and PMP populations decline, while hub cells rise, reestablishing β-cell functionality.

Clearly, the interplay between β-cell dedifferentiation, nascent β-cell maturation, and β-cell subtype identity in diabetic-obesity needs to be clarified. SM/J mice are a useful model because they allow for appropriate comparisons across diabetic-obese, nondiabetic-obese, and nondiabetic-lean populations. Future studies interrogating these differences will provide insight into the physiological mechanisms that allow β-cell functionality to be maintained and improved in the context of obesity.

## Acknowledgements

### Author Contributions

HAL and MAM designed the experiments. MAM, CC, HS, and JPW performed mouse phenotyping and ELISAs. MAM, JMV, and MK performed histological assays and analyses. MAM, JWH, and CLP performed experiments on isolated islets. MAM wrote the manuscript. All authors edited and approved the final draft.

### Funding

This work was supported by the Washington University Department of Genetics, the Diabetes Research Center at Washington University (P30DK020579), the NIH NIDDK (K01 DK095003) to HAL and NIH NIGM (T32 GM007067) and NIH NIDDK (T32 DK108742) to MAM

The authors declare no conflicts of interest.

