## Supplemental Figures and Table for "Spontaneous restoration of functional β-cell mass in obese SM/J mice"

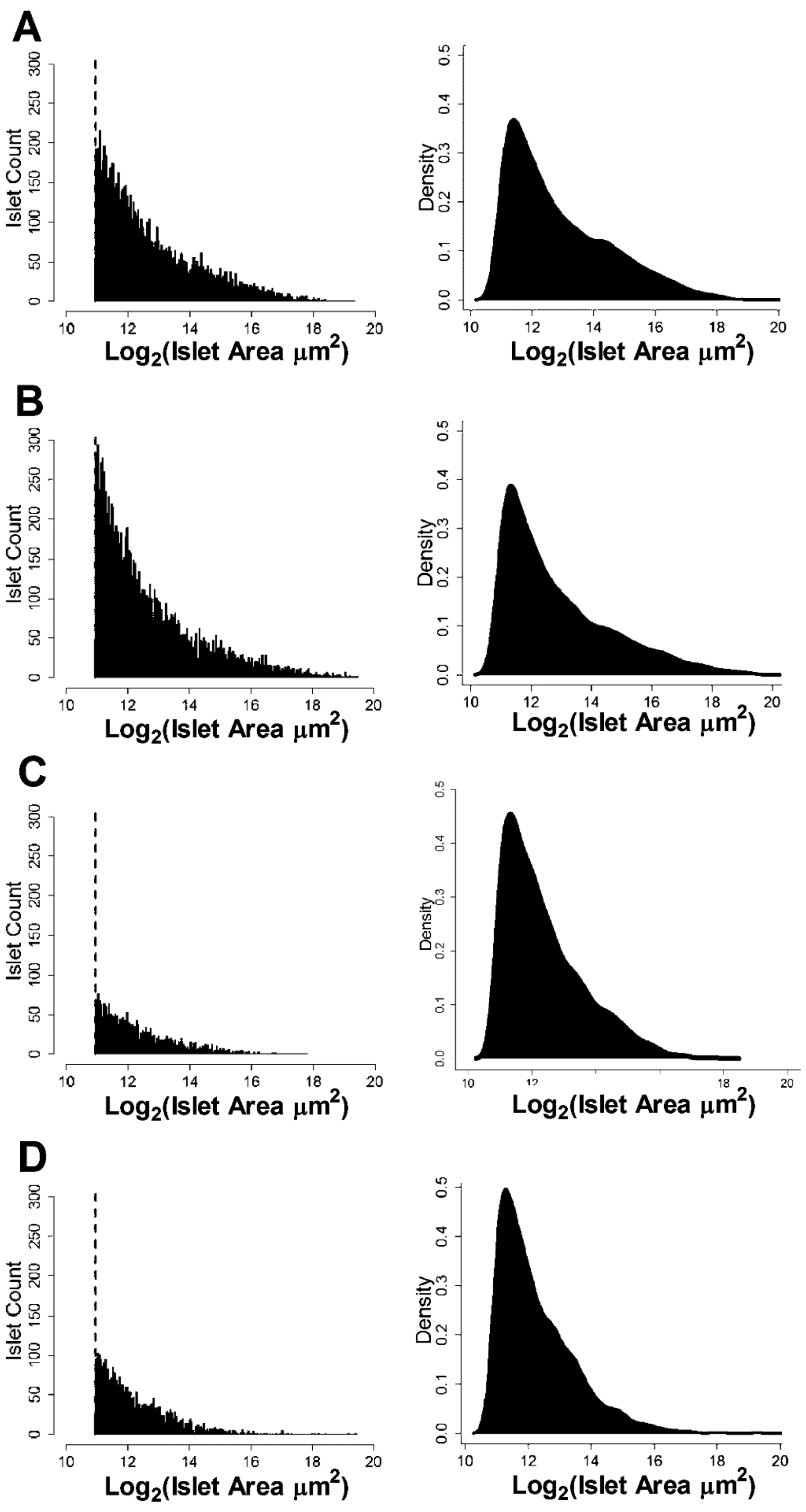

A
B
C
D
Islet Count
Density
Log2(Islet Area μm2)

**Supplemental Figure 1**. Islet area histogram (left) and density plot (right) for each cohort, 20-week high fat **(A)**, 30-week high fat **(B)**, 20-week low fat **(C)**, and 30-week low fat **(D)**. Dashed line shows islet size threshold. N= 2,333-10,380 islets per cohort.

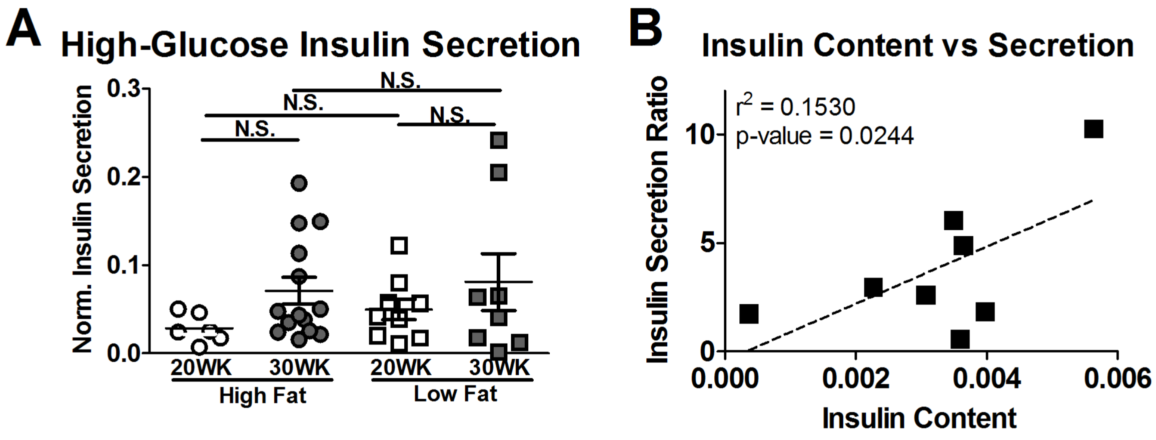


**Supplemental Figure 2**. Normalized insulin secretion in response to elevated (11 mM) glucose **(A)**. Islet insulin content vs glucose-stimulated insulin secretion for low fat-fed mice **(B)**. A, N =5-14 individuals per cohort. B, N = 8. N.S. – Not significant

**Supplemental Table 1**. Cohort metrics for islet mass quantification.

| Cohort | Individuals | Female /Male Ratio | Slides | Total Islets | Total Islet Area (µm²) | Avg Iset Area (µm²) | Islets/Slide | Area/Slide | Total Tissue Area (µm²) | Islet Area/Tissue Area |
|---|---|---|---|---|---|---|---|---|---|---|
| 20 HF | 16 | 0.5 | 62 | 8,320 | 132,564,412 | 15,933 | 134.2 | 2,138,136 | 496,461,183 | 0.27 |
| 30 HF | 16 | 0.5 | 64 | 10,380 | 194,565,474 | 18,744 | 162.2 | 3,040,086 | 512,715,812 | 0.38 |
| 20 LF | 12 | 0.5 | 48 | 2,333 | 19,956,617 | 8,554 | 48.6 | 415,763 | 176,863,435 | 0.11 |
| 30 LF | 11 | 0.64 | 40 | 3,141 | 28,581,981 | 9,100 | 78.5 | 714,550 | 198,371,139 | 0.14 |
